# Photosynthesis under very high oxygen concentrations in dense microbial mats and biofilms

**DOI:** 10.1101/335299

**Authors:** Dirk de Beer, Volker Meyer, Judith Klatt, Tong Li

**Author notes:** Address correspondence to Dirk de Beer.

## Abstract

Using microsensors O_2_ concentrations were measured in photosynthetically active microbial mats of up to 3 mM, corresponding to a partial pressure of 3 bar. This could damage mats by internal gas formation, and be inhibitory by formation of reactive oxygen species (ROS) and reduced effectivity of RuBisCo. The reliability of the electrochemical microsensors was checked by creating elevated O_2_ concentrations in a water volume placed inside a pressure tank. A microsensor mounted with the tip in the gassed water bath showed a response linearly proportional to 5.5 mM corresponding to 4 bar pure O_2_ pressure. After release of the pressure the O_2_ concentration reduced quickly to 2.5 mM, then stabilized and subsequently reduced slowly over 14 hours to approximately 2 mM. We concluded that the very high O_2_ concentrations measured in phototrophic microbial mats are real and O_2_ oversaturation in mats is a stable phenomenon. As consequence of high O_2_ concentrations, net production of H_2_O_2_ occurred. The accumulation was, however, limited to the respiratory zone under the photosynthetic layer. Despite the high gas pressure inside mats, no disruption of the mat structure was apparent by bubble formation inside the mats,and bubbles were only observed at mat surfaces. Additions of H_2_O_2_ to high concentrations in the water column were efficiently removed in the photosynthetically active zone. As the removal rate was linearly proportional to the H_2_O_2_ influx, this removal occurred possibly not enzymatically but by abiotic processes. Phototrophic microorganisms can produce O_2_ at high rates under strongly elevated O_2_ levels, despite the decreased efficiency due to the unfavorable kinetics of RuBisCo and energy costs for protection. Under non-limiting light conditions, this apparent dilemma is, however, not disadvantageous.

**Importance:** Biofilms are often used in photobioreactors for production of biomass, food or specialty chemistry. Photosynthesis rates can be limited by high O_2_ levels or high O_2_/CO_2_ ratios which are especially enhanced in biofilms and mats, due to mass transfer limitations. High O_2_ may lead to reactive O_2_ species (ROS) and reduce the efficiency of RuBisCo. Moreover, gas formation may destabilize their structure. Here we show that extremely high levels of O_2_ are possible in mats and biofilms without ebullition, and while maintaining very high photosynthetic activity.

## Introduction

In lakes and coastal waters O_2_ levels modestly higher than air-saturation is commonly observed (1, 2). In phototrophic microbial mats and microphytobenthos much higher O_2_ levels develop due to mass transfer resistance, and indeed 2-5 times air saturation is commonly observed with electrochemical O_2_ microsensors (3-6). High O_2_ levels, especially under low CO_2_, can enhance the oxygenase activity by ribulose-1,5-bisphosphate carboxylase-oxygenase (RuBisCo) leading to a reduction of net photosynthesis (7). This would preclude the formation of high levels of O_2_ by phototrophs, and be especially inhibitory for photosynthesis in mats and biofilms, where due to mass transfer resistance O_2_ can accumulate and CO_2_ can be depleted. However, detailed studies on hypersaline intact mats showed no effect of elevated O_2_ concentrations on net- and gross photosynthesis (8, 9). These studies reported O_2_ levels of up to 5 times air saturation, equaling 1 bar of pure O_2_. This is remarkable considering the potential cell damage by extreme O_2_ levels. O_2_ in its ground state is rather unreactive unless catalyzed, however, organisms in oxic environments run the risk of being damaged by formed ROS. O_2_-derived ROS comprise superoxide, hydrogen peroxide (H_2_O_2_), and hydroxyl radicals(10). ROS can be formed by O_2_ reacting with cell components that are brought in high energy state in photosynthesis(11) or respiration(12). These activated cell components will more readily produce ROS when the O_2_ is in high concentration, as occurs in illuminated phototrophic mats. During photosynthesis light-activated reaction centers in photosystem I and II (PSI and PSII) can react with O_2_ to generate ROS. Particularly under high illumination, chlorophyll-a (Chl-a) triplets accumulate, that react with O_2_ resulting in the formation of singlet O_2_ in PSII and superoxide anion radicals in PSI (Mehler reaction). During respiration activated quinones in the electron transport chain and the terminal electron donor, cytochrome-c oxidase, are the sources of ROS formation. A well-known defense mechanism is the combination of superoxide dismutase that converts superoxide into H_2_O_2_ and catalase that degrades H_2_O_2_ to water and O_2_. For H_2_O_2_ microsensors are available. A last issue is the potential for mat disruption by gas formation. Indeed, often bubbles are observed and released from these photosynthetic communities, which indicates internal oversaturation, O_2_ partial pressures exceeding atmospheric pressure (9, 13). Especially under conditions where DIC is not limiting, as occurs in alkaline lakes or in phototrophic bioreactors with purposefully elevated pH, extremely high O_2_ levels strongly exceeding partial pressures of 1 bar pure O_2_ are theoretically well possible inside biofilms. However, reports on oversaturation are rare, possibly, such observations were considered artifacts. We report extremely high O_2_ levels in biofilms and mats of up to 3 times pure O_2_ saturation, so 15 times air-saturation. We tested the reliability of our measuring technique, the electrochemical O_2_ microsensors, under extremely high O_2_ tension. Furthermore, we measured the magnitude of the possible oversaturation in water, and determined the stability of the oversaturation. Also, we assessed the spatial distribution of H_2_O_2_ in highly active mats to assess community defense mechanisms against ROS.

## Results

In different biological systems, mats and biofilms, we measured significant oversaturations of O_2_ upon illumination (Fig. 2). At the surface of all mats bubbles were formed during illumination, but no escaping bubbles from within the mats occurred and no evidence of structural damage by bubbles was obvious. In the microbial mat from Solar Lake the highest measured O_2_ concentration was approximately 1.1 mol m^−3^, which was over 7 times air-saturation (Fig. 2A). In the microbial mats from the alkaline lake Boitano (Fig. 2B), we measured O_2_ maxima of 2.2 mol m^−3^, which is 9 times above air-saturation. The overlaying water column had an oversaturation in O_2_, due to the limited water-air exchange. The biofilms grown in alkaline saline water had the highest volumetric activity (Fig. 2C), with an O_2_ peak of 2.3 mol m^−3^.

It should be noted that due to the different salinities and temperatures, the air-saturated O_2_ concentration was different in each experiment. In the Solar Lake water O_2_ was at air-saturation 0.146 mol m^−3^, in the water of Lake Boitano 0.26 mol m^−3^ and in the biofilm experiment 0.16 mol m^−3^.

To test the response of sensors above pure oxygen saturation, a sensor was placed in the pressure tank (Fig. 1) and the O_2_ level in the water was increased to 5.5 mol m^−3^ (Fig. 3), by stepwise increasing the pressure to 4 bar during bubbling the water. The resulting calibration was perfectly linear (Fig. 4). After the sensor calibration the tank was vented and the pressure was released to atmospheric level in approximately 10 minutes. The O_2_ concentration quickly (in one hour) reduced from 5.5 mol m^−3^ to 2.5 mol m^−3^ and then reduced very slowly in the subsequent 14 hours to 2 mol m^−3^. After that the tank and water volume was ventilated by air, and the signal of the sensor returned to that of air saturation. To assess the effect of high O_2_ on ROS development we measured H_2_O_2_ profiles as function of illumination (Fig. 5A and B). In illuminated mats, O_2_ did not reach oversaturation, but approached pure oxygen saturation. The H_2_O_2_ sensor measured a background signal, corresponding to 0.5-1 µM H_2_O_2_, that disappeared in the permanently anoxic zones in the mats. Deep in the mats (below 5 mm in the light, below 3 mm in the dark) the signal went up sharply, probably due to sulfide interference (not shown). H_2_O_2_ showed a maximum in the respiratory zone of the mats, clearly below the photosynthesis driven O_2_ maximum, and this peak was elevated in the light (Fig. 5). Upon addition of increasing concentrations to the overlaying water H_2_O_2_ did not penetrate to below the photosynthetic layer (Fig. 6A). All influxing H_2_O_2_ was completely consumed at 1.1-1.3 mm depth, independent off the concentration in the water column. Consequently, the conversion rate in the mats (calculated from the interfacial flux divided by the penetration depth) was linearly proportional to the concentration in the water column (Fig. 6B). The H_2_O_2_ additions did not reduce the oxygen production upon illumination (Fig. 6A).

**Fig. 1.**
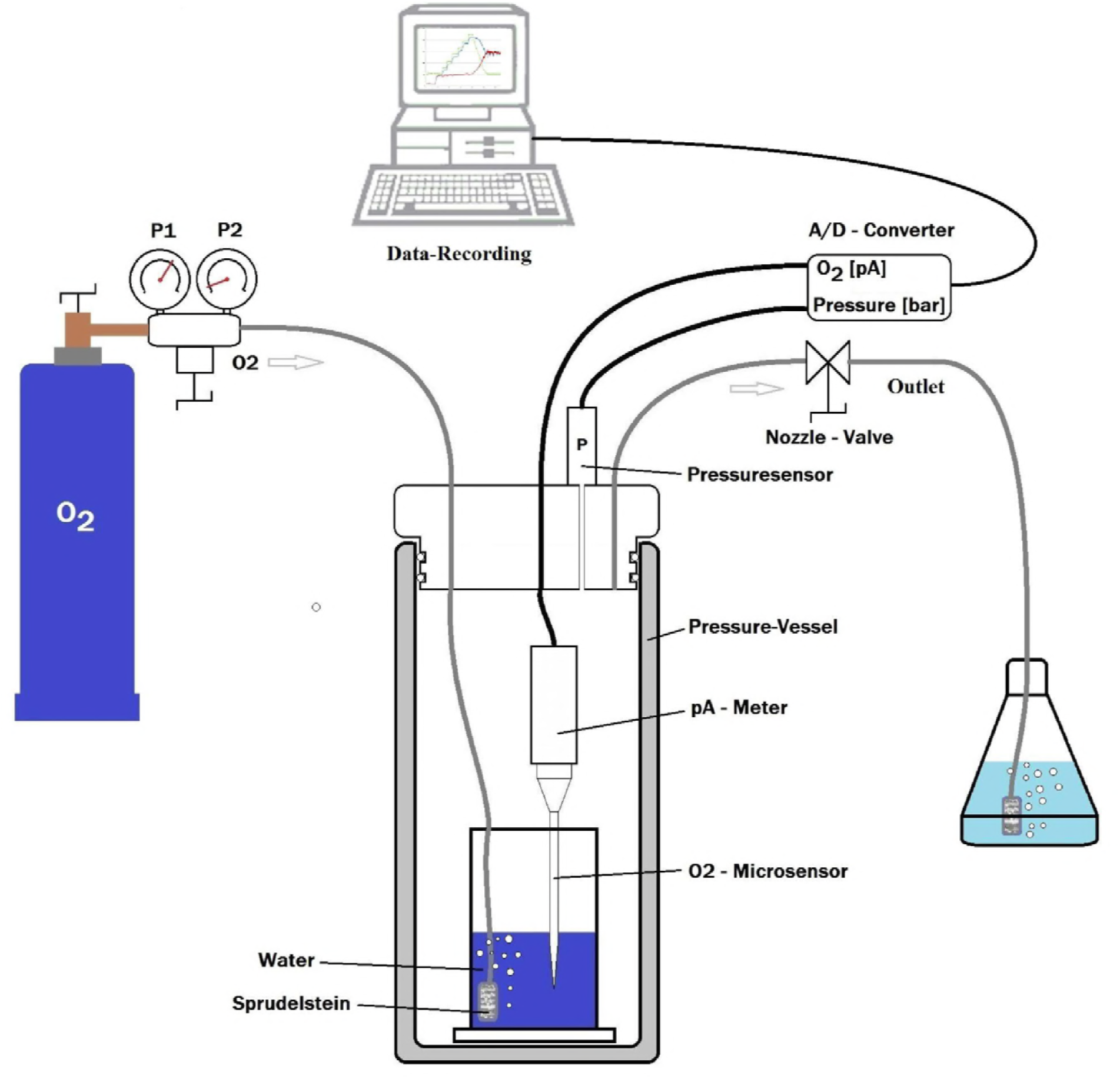
Scheme of the pressure tank for experiments with O_2_ pressures up to 5 bar.

**Fig 2.**
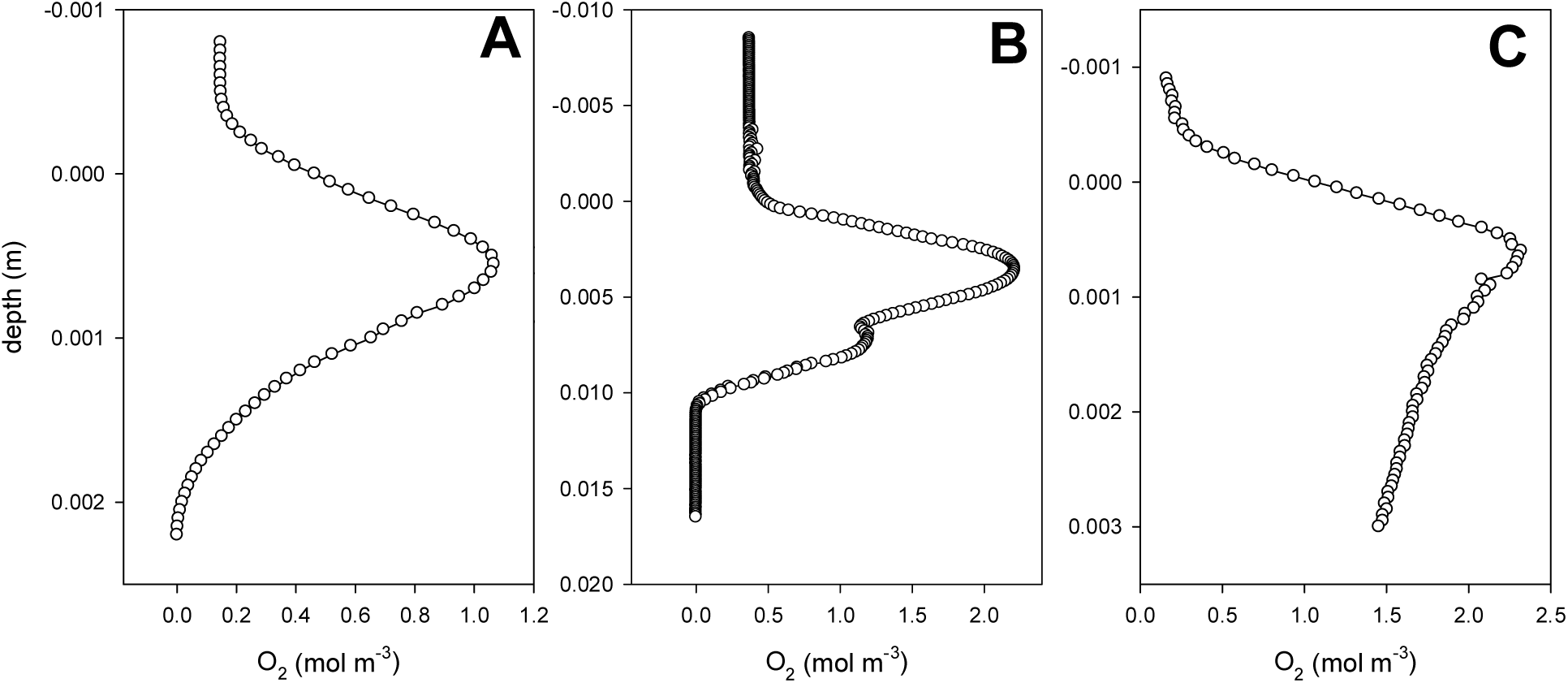
O_2_ profiles measured in a microbial mat from Solar Lake, Sinai (A), in a mat from Lake Boitano, Canada (B), and a laboratory biofilm grown at high pH and 1 M DIC (C).

**Fig. 3.**
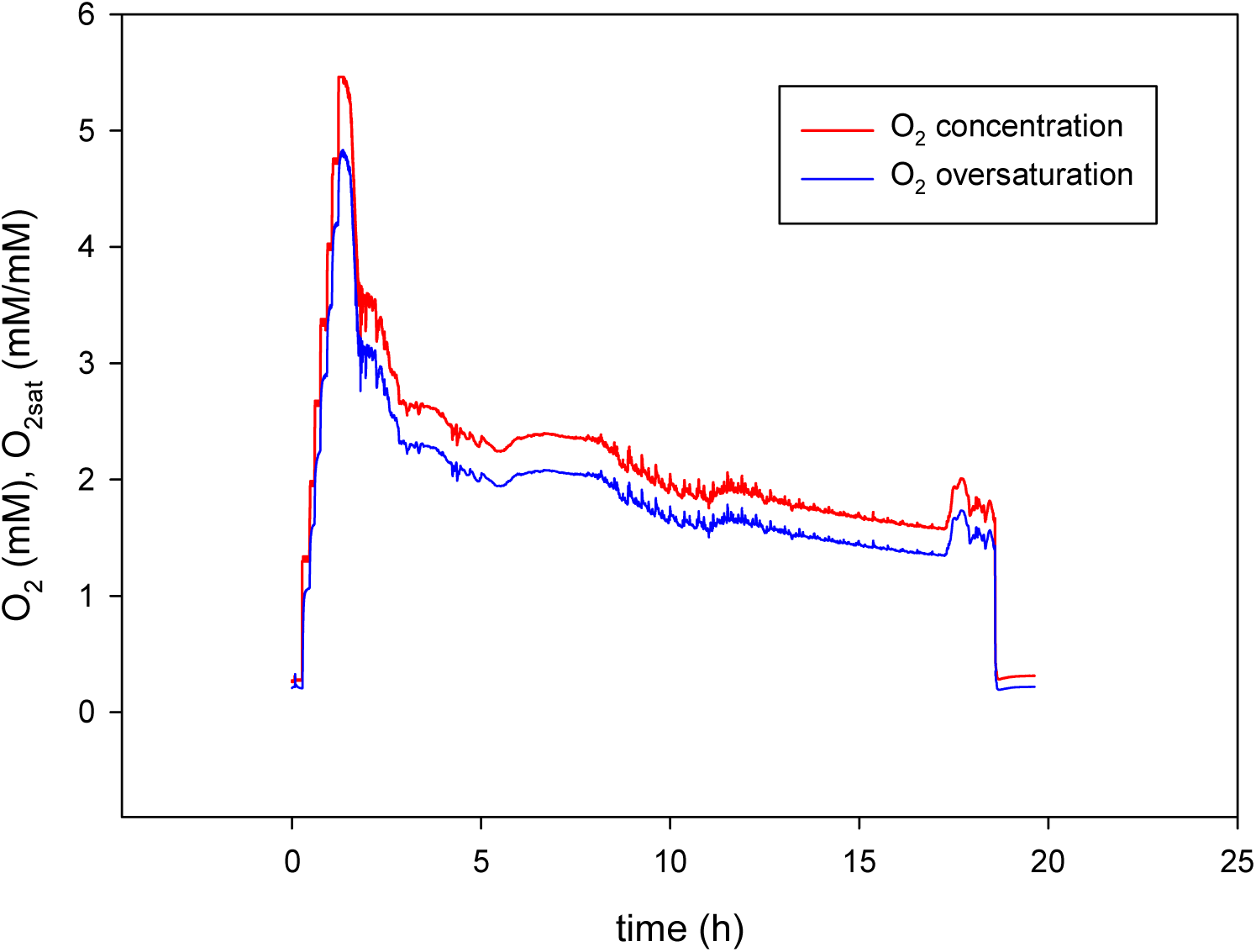
O_2_ concentrations during the experiment in the pressure tank, and the oversaturation level calculated by dividing the measured concentration with the saturation level at 1 bar of pure O_2_. In the first 1.5 hours the O_2_ tension was stepwise increased to record the response of the microsensor to extreme O_2_ concentrations. Subsequently, the pressure was released in 10 minutes to 1 bar, and the O_2_ dynamics of the further degassing was followed overnight, in a locked room. At 18 hours the tank was ventilated by air.

**Fig. 4.**
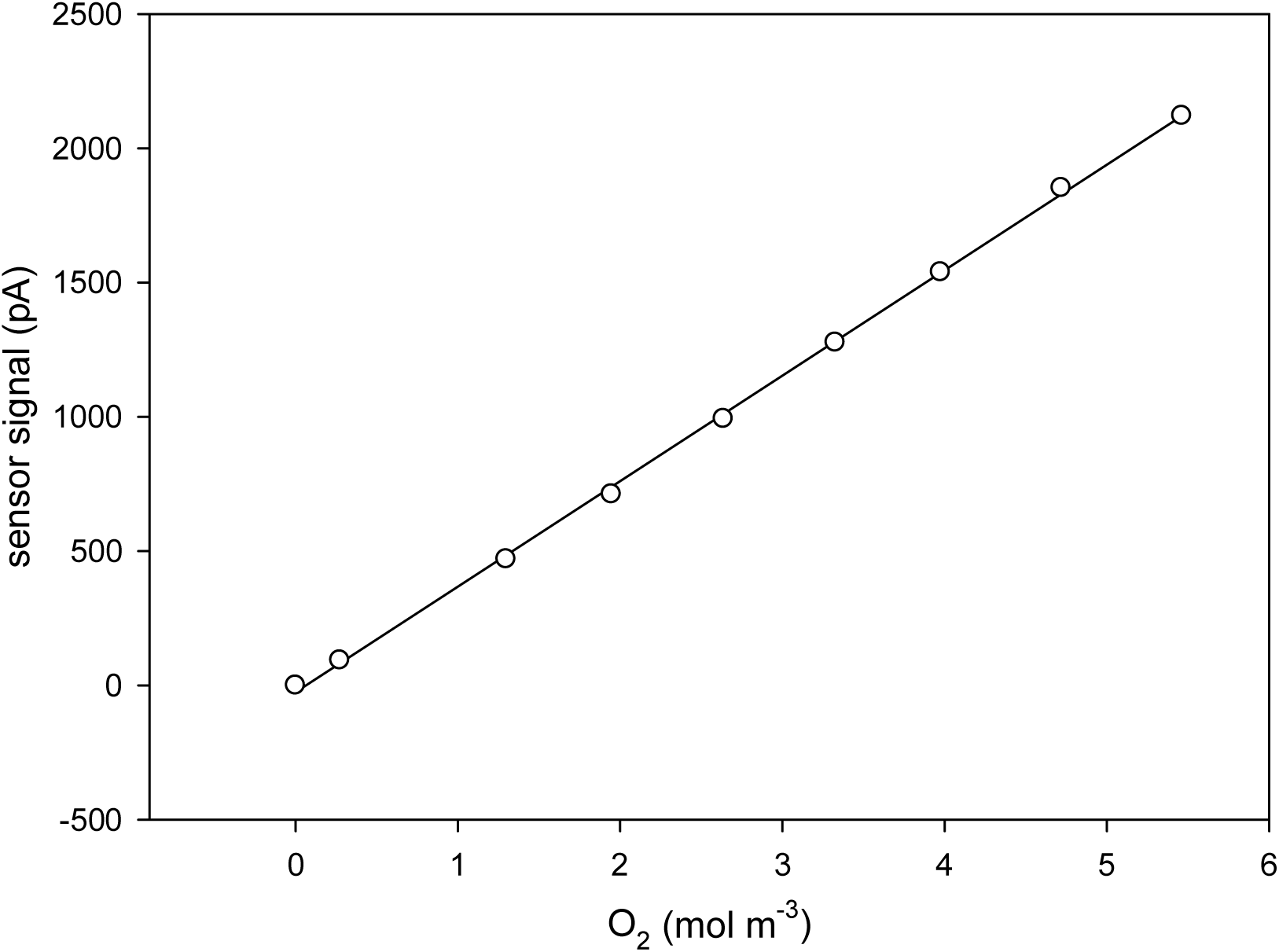
The calibration line of an O2 microelectrode to very high concentrations. The line is the linear regression (R^2^=0.999).

**Fig. 5.**
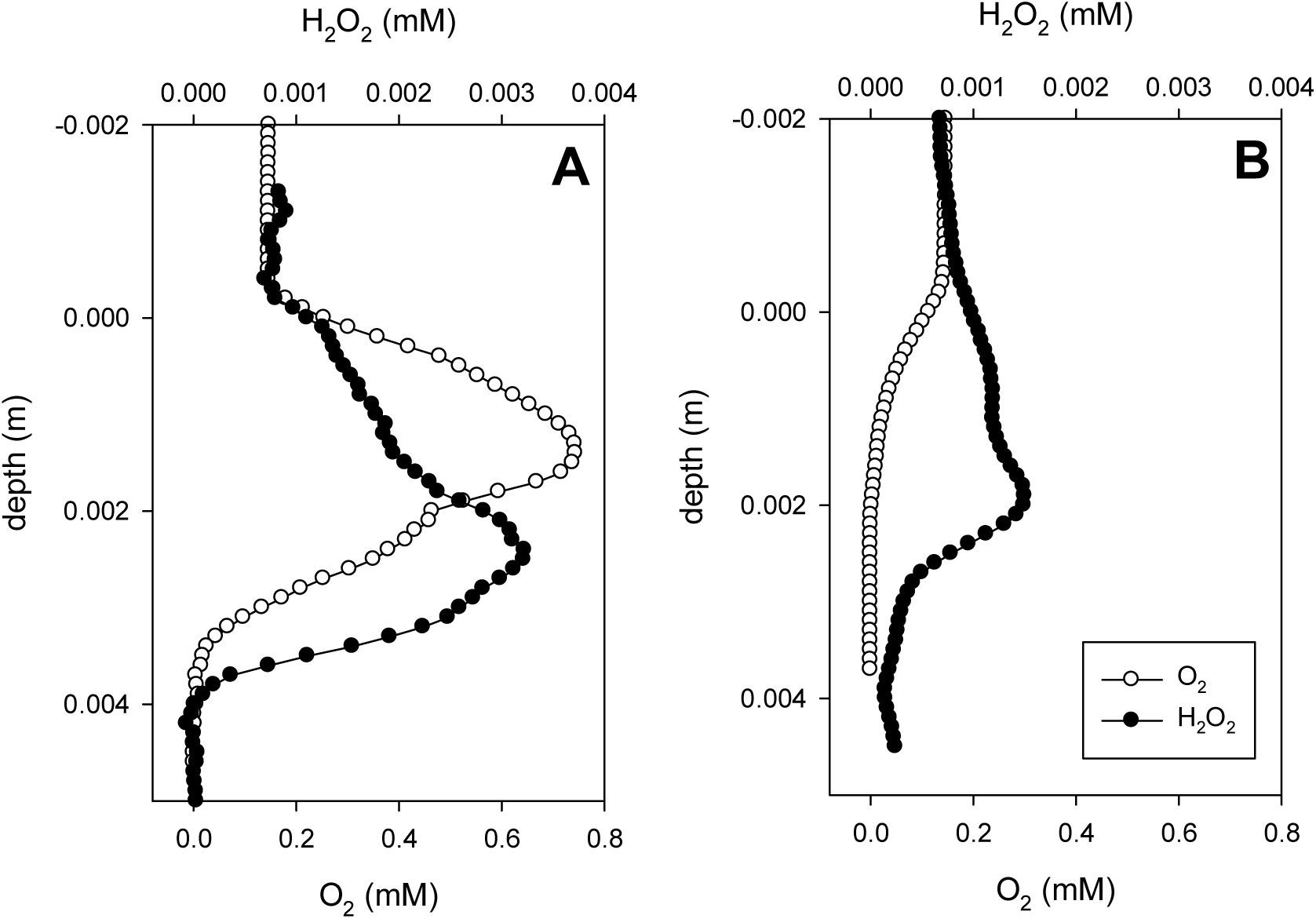
O_2_and H_2_O_2_microprofiles measured in an illuminated (A) and dark-incubated (B) cyanobacterial mat.

**Fig. 6.**
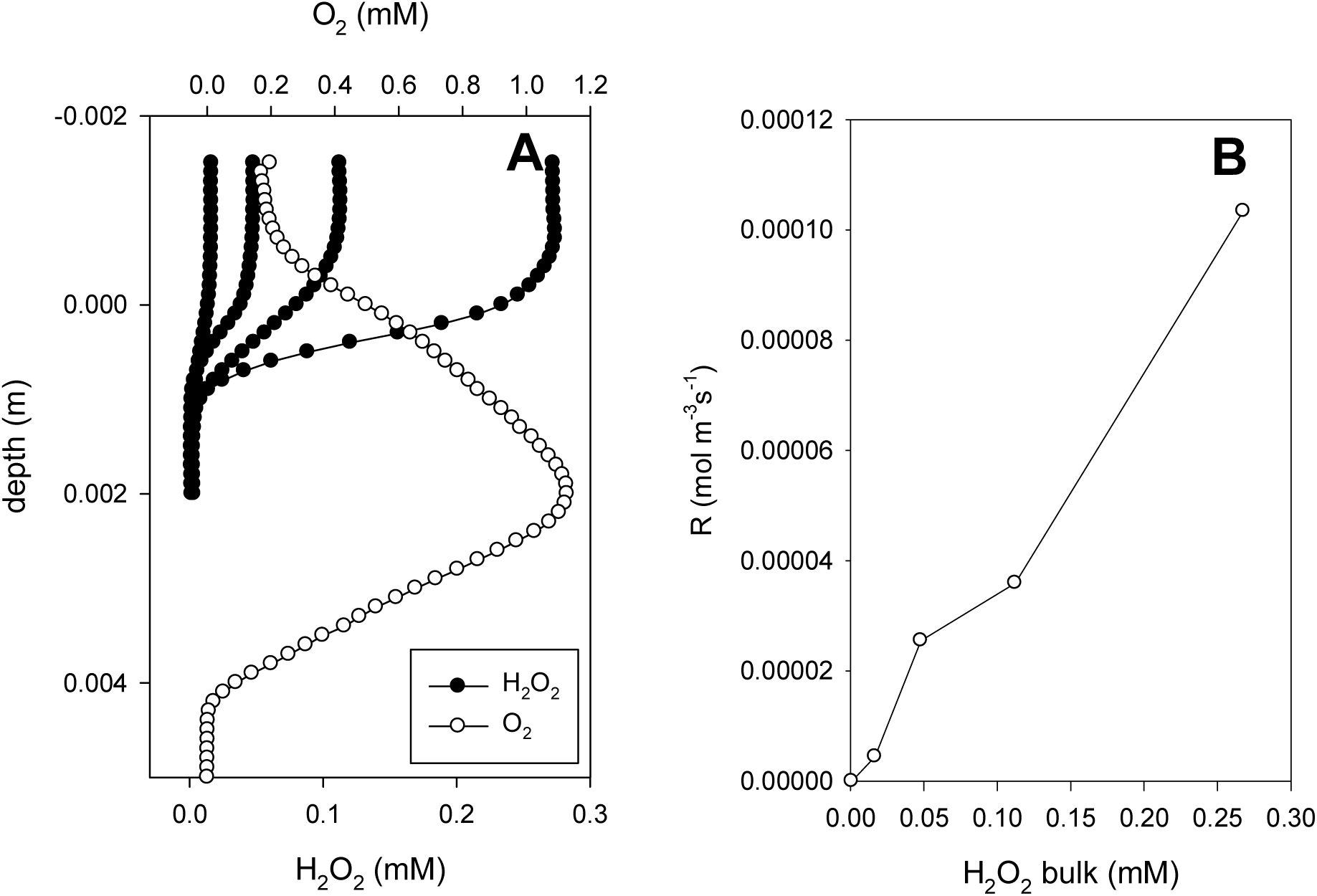
H_2_O_2_ profiles measured in a cyanobacterial mat with increasing additions of H_2_O_2_ (17, 48, 110 and 272 µM) to the overlying water column (Fig. 6A). From the penetration depths and the interfacial fluxes the volumetric H_2_O_2_ consumption rates were calculated (Fig. 6B). After the last H_2_O_2_ addition illumination was switched on and, after reaching steady state, an O_2_ profile was measured, showing high photosynthetic activity and no damage by H_2_O_2_.

## Discussion

O_2_ sensors are usually calibrated in O_2_ free and air saturated water, and occasionally in pure O_2_. To test whether observations of higher O_2_ levels in mats are real, we needed to extend the calibration range to above 1 bar pure O_2_. The microsensor worked perfectly in 5 bar partial pressure O_2_. After the exposure to extreme O_2_ levels the sensor response was unchanged. Thus very high levels of O_2_ can reliably be measured. Oversaturation can remain stable for at least 12 hours (Fig. 3), much longer than the diffusion times (t), calculated using t=x^2^/(2D), where x is the distance and D the diffusion coefficient. The O_2_ maxima in the mats occurred at 0.001-0.005 m depth (Fig. 2), resulting in diffusion times of 4 minutes to 1.5 hour. Consequently, extreme O_2_ oversaturation in actively photosynthesizing mats and biofilms is indeed very well possible.

We then considered whether such high O_2_ concentration, above 1 bar partial pressure, can occur without leading to instant ebullition. Oversaturation and bubble formation in liquids is studied since decades, and appears highly complex(14). The pressure inside bubbles is inversely related to their size, as surface tension increases with decreasing bubble diameter. The smallest bubbles are subjected to extremely high (up to 260 atm) surface tensions (15). The ensuing very high internal pressure ‘squeezes’ gasses out of the bubbles into solution. Thus small bubbles are unstable and tend to dissolve increasingly fast while shrinking. Conversely, very high gas concentrations are needed to overcome the surface tension and to create a bubble. Once a bubble is formed and expands the surface tension goes down and its internal pressure decreases, and it thus absorbs increasingly fast gas from the liquid. There is a critical size below which bubbles dissolve faster than they can grow, even in oversaturated medium. Above the critical size they will grow faster than they dissolve (16).

Thus in clean water, the site of O_2_ production (e.g. an electrode) O_2_ levels must be ∼300 times higher than saturation, close to 2.5 M, before formation of bubbles can occur (17). This concentration is needed to overcome the surface tension of 260 atm of the initial bubbles. Such partial pressures seem beyond physiological ranges, hence bubble formation by photosynthesis in biological systems like mats and sediments requires an additional mechanism. Formation of bubbles can be stimulated by nucleation on heterogeneous surfaces nucleation (18). Nucleation sites are hydrophobic micro-areas on surfaces, which stabilize microbubbles (<1 µm) and even allow their growth to above the critical size above which bubbles grow faster than they dissolve(14). The surrounding solution must be oversaturated, the local concentrations depend on the transport rate of gas to the bubbles. Hydrophobic microsites are indeed abundant on cell surfaces(19, 20). Their density and accessibility is critical for bubble formation(18). Degassing of fluids is a function of the surface tension of the liquid, size and internal pressure of bubbles, the viscosity of the liquid and the diffusion coefficient of the gas in this liquid, the release rate of the bubble to the surface and foam building leading to transfer resistance between liquid and air. Ebullition is thus a highly complex phenomenon, but strong oversaturation is very well possible with a degree depending on the density of nucleation sites, their surface properties and mass transfer phenomena. We suspect that potential nucleation sites in phototrophic mats and biofilms are somehow shielded, so that structural damage by internal bubble formation is prevented.

As supersaturation in mats is real and does not necessarily lead to bubbling, the exposed organisms have to deal with the physiological consequences of extreme O_2_ levels. The physiology of phototrophs and the whole community can be affected in various ways, via substrate stimulation or product inhibition phenomena. Most prominently, the community is expected to suffer from exposure to ROS. Indeed, we observed net production of H_2_O_2_ in mats especially under high photosynthesis rates, but only by respiratory activity below the photic zone. Both photosynthesis and respiration are sources for H_2_O_2_. Apparently, especially the phototrophs have an efficient protection mechanism against H_2_O_2_ by active degradation of the compound. The power of this protection is demonstrated by the experiment where we enhanced the medium concentration to very high levels. The consumption of H_2_O_2_ in the photosynthetic active zone of the mats was first-order suggesting that the degradation was not enzymatically mediated. If H_2_O_2_ were converted enzymatically, its affinity to H_2_O_2_ must be very low, which is unlikely for a protective mechanism. The H_2_O_2_ penetrated to 1.2 mm depth, independent of the concentration. At that location, possibly pools of labile reduced substances are stored by the phototrophs, as protection against ROS. The phototrophic activity of the mats was not damaged by the exposure to extreme H_2_O_2_ levels. In the deeper zones of the mats where O_2_ consumption dominates, protection against H_2_O_2_ appears to be less active, as here the H_2_O_2_ peak was enhanced by illumination, due to increased respiration, driven by higher fluxes of photosynthetically produced O_2_.

Despite the remarkable protection again ROS, photosynthetic organisms still have to cope with potentially decreased carbon fixation efficiency in the presence of high O_2_. This is because of increased O_2_ consumption by photorespiration, the oxygenase reaction of RuBisCo under formation of phosphoglycolate (21). It is well documented that RuBisCo is sensitive to O_2_, as the oxygenase activity is controlled by the O_2_/CO_2_ ratio(22). This effect is demonstrated in reconstituted enzyme systems and spinach chloroplasts, but is much less obvious in intact microbial cells. Aquatic microalgae have a highly effective carbon concentrating mechanism (CCM) (7, 23, 24), that elevates the intracellular CO_2_ concentrations by several orders of magnitude and thereby effectively blocks the oxygenase reaction (25). Moreover, the affinity ratio (ratios of K_m_ for CO_2_ and O_2_) of RuBisCo from microalgae is lower than of plants (26). Hence also photorespiration is unlikely to be strongly affected by extreme O_2_ levels, even when under 3 mM of O_2_ in the cell surroundings an energy expensive CCM is needed.

The consequences of high O_2_, enhanced ROS and enhanced oxygenase activity of RuBisCo, are countered with considerable energy cost. For example, a large part of the intracellularly accumulated dissolved inorganic carbon (DIC) leaks out again as CO_2_ permeates easily through membranes(27, 28). While all the protective leaks and photorespiration are reducing photosynthetic efficiency, it seems not to be of a disadvantage, as ROS mainly develops when photosynthesis is not light limited. Protection is clearly more important than energy efficiency under such conditions. This is also manifest in the fact that the highly active mechanisms to remove superoxide and hydrogen peroxide can make cyanobacterial mats sinks for these compounds: the mats are able to remove much higher H_2_O_2_ fluxes than required for protection against their internal production.

We are still only scratching the surface of the consequences of high O_2_ and ROS, and important regulatory functions are likely to be discovered. Further experiments with the pressure tank will be done to better relate O_2_ and DIC levels to photosynthesis rates, energy efficiency and ROS production. Furthermore, due to the enormous partial pressure of O_2_ that naturally develops in cyanobacterial mats, these are ideal to study effects of high O_2_ on phototrophs and other inhabitants.

## Materials and methods

Microbial mats were collected from Solar Lake (Sinai, Egypt, 29°25’20.66“N 34°49’47.66”E) in 1997, brought to the laboratory in Bremen, Germany, and stored at 27°C in artificial seawater with ambient salinity (10.5%), pH 8.2, under irradiance of 400 µmol photons m^−2^s^−1^, with 12-12 hour diel cycle. The measurements were done within two weeks after sampling, under the same conditions, while the water column was bubbled with air.

Microbial mats were sampled from Lake Boitano (British Columbia, Canada, 51°56’58.02“N 122° 7’45.91”W) in June 2017, brought to the laboratory in Calgary and stored at room temperature under irradiance of 136 µmol photons m^−2^s^−1^, with 12-12 hour diel cycle. The salinity was 0.9%, the pH of the water was 10.5. The measurements were done in two days after sampling at 300 µmol photons m^−2^s^−1^. An airjet was blown over the water column to maintain a gentle current.

The biofilms were grown from an inoculum obtained from hyperalkaline lake mats (29) in artificial hyperalkaline medium with 85 g L^−1^ sodium bicarbonate on agar plate submerged at 2 cm depth. The irradiance during cultivation was 100 µmol photons m^−2^s^−1^, with 12-12 hour diel cycle. The salinity was 8.5%, the pH was 8.9 the temperature 20°C. The microsensor measurement was performed 5 days after inoculation. Light intensity during measurement was 100 µmol photons m^−2^s^−1^, mixing was provided by a gentle air current above the water surface.

Mats were collected from the d’Es Trenc salina (Mallorca, Spain, 39°20’44.00“N 3° 0’1.73”E) and were maintained at 11% salinity under 300 µmol photons m^−2^s^−1^, with 12-12 hour diel cycle, at 20°C. Measurements on the effect of H_2_O_2_ were performed 4 months after sampling, mixing was provided by a gentle air current above the water surface.

O_2_ microsensors with a guard cathode and a tip diameter of 10 µm were prepared, calibrated and used as described previously (30, 31). Single anode H_2_O_2_ electrodes were prepared as described previously (32). Shortly, glass-coated Pt wires were tapered by etching at 2.5 V in saturated KCN and coated with glass in a hot Pt loop, opened at a diameter of 10 µm and reheated to reseal the glass to the Pt wire. Their tips were recessed to 20 µm from the tip, at 2.5 V in saturated KCN. The recesses were filled with porous Pt by electroplating in PtCl_6_ (2% in 1M HCl) under microscopic observation until the recesses were filled after approximately 5 minutes. These electrodes were coated with cross-linked bovine serum albumin (BSA). For this 1 ml 10% BSA in 50 mM phosphate buffer pH 7.3 was vortexed quickly with 10 µL 50% glutaraldehyde, a drop of the mixture was brought in a Pasteur pipette that was made slightly thinner in the tip, and under microscopic guidance the electrode was moved in and out of the mixture until the BSA became syrupy and a thin film covered the electrode. This was dried in air overnight after which a ceramic-like coating developed. A potential was applied of +700 mV against a Ag/AgCl reference, and the sensor was stabilized for 10 minutes in the saline incubation medium and calibrated by adding aliquots of H_2_O_2_ from a 3% stock that was stabilized by phosphoric acid. The sensor was highly sensitive to sulfide resulting in an elevated signal, and entirely insensitive to O_2_. The diffusion coefficient of H_2_O_2_ in the hypersaline mats was 5.25 x 10^−10^ m^2^ s^−1^, obtained from (33), corrected for salinity and the mat matrix(3). A pressure tank was used to investigate the response of the sensor to high O_2_ levels (Fig. 1). The tank was made from stainless steel and had a volume of 11.3 L. The sensor was mounted to the pressure-resistant housing of the amplifier. The amplifier provided the potential of −0.8 V and measured the current of the microsensor. The amplifier was connected with a cable through the lid of the pressure tank to a laptop with a program for recording the microsensor signals. The sensor tip was positioned in a beaker with water. The water could be bubbled with pure O_2_, supplied from a tank with O_2_ gas. All gas in the tank first passed the water beaker with the O_2_ microsensor. The pressure was regulated by the second stage of the regulator and the pressure was recorded, with a pressure sensor, simultaneously with the microsensor signals. A release valve was used to regulate the throughput of pure O_2_ gas. Initially the water was air-saturated.

During the experiment the gas content of the cylinder was first exchanged by pure O_2_ at 1 bar. Then the pressure was stepwise increased, while bubbling continued. Each step was performed until the sensor signal was stable, which took ca 15 minutes. After the highest pressure test of the sensor, the pressure was released in 20 minutes and the cylinder was left untouched overnight, while the recording of sensor signals continued.

## Acknowledgements

We thank our technical staff for supplying the microsensors. This research received no specific grant from any funding agency in the public, commercial, or not-for-profit sectors.

